# Systematic analysis of intracellular replication competence of mammalian, avian, and reptilian kolmioviruses

**DOI:** 10.64898/2026.09.08.748734

**Authors:** Mai Kishimoto, Hiroki Furukawa, Akari Sono, Ryosuke Fujimura, Mina Isomichi, Masashi Iwamoto, Koichi Watashi, Ryusei Kuwata, Masayuki Horie

**Author notes:** Corresponding authors Mai Kishimoto, DVM, PhD, Masayuki Horie, DVM, PhD. These authors contributed equally to this work (Mai Kishimoto, Hiroki Furukawa).

## Abstract

Kolmioviruses (KoVs) are circular, single-stranded RNA viruses whose evolution may have involved host switching between vertebrate classes. However, the host specificity of their intracellular replication remains poorly understood. Here, we examined six mammalian, one avian, and one reptilian KoVs using cell lines derived from the same three vertebrate classes. We transfected cells with plasmids encoding 1.2× genome-length KoV sequences and assessed persistent replication by monitoring delta antigen-positive cells for up to 18 days. Four mammalian KoVs replicated persistently only in mammalian cell lines, whereas avian and reptilian KoVs replicated persistently only in both avian and reptilian cell lines. By contrast, the remaining two mammalian KoVs, nested within the clade whose deepest lineages are the avian and reptilian KoVs, replicated in cell lines from all three vertebrate classes. The observed patterns of persistent replication are concordant with the phylogenetic grouping of the KoVs. Our findings support a possible evolutionary scenario in which the common ancestor of the two mammalian KoVs in the avian–reptilian clade acquired compatibility with mammalian cells while retaining the capacity to replicate in avian and reptilian cells, contributing to the transition of this lineage to mammalian hosts.

## Main Text

Kolmioviruses (KoVs; family *Kolmioviridae*, realm *Ribozyviria*) are small, circular, single-stranded RNA viruses (1). Hepatitis D virus (HDV), first identified in humans in 1977 (2), remained the only known member of this group for several decades. Since 2018, however, diverse HDV-like viruses have been identified in a wide range of animal species (3–8), leading to the establishment of the family *Kolmioviridae* (1). Molecular evolutionary analyses suggest that host switching has played an important role in the evolution of KoVs. Notably, mammalian KoVs consistently fall into two distinct phylogenetic lineages (7–9): one comprises only mammalian KoVs, whereas the other has avian and reptilian KoVs as its deepest lineages, with the mammalian KoVs nested among them. This nested position raises the possibility that some mammalian KoVs originated through host switching between vertebrate classes. However, the host ranges of individual KoVs remain poorly understood, making this possibility difficult to assess.

KoVs require envelope proteins supplied by helper viruses to form infectious particles. Their host specificity is therefore determined by at least two major barriers: helper virus-mediated entry into host cells and subsequent replication of KoV genomes within those cells. Although the ability to use diverse helper viruses may facilitate transmission to new hosts, successful host switching also requires KoVs to replicate in cells of the recipient host. Experimental studies have shown that several KoVs can use viruses from different families as helpers (10, 11) and that some can replicate in cultured cells derived from species other than their known hosts (7, 12, 13). However, these studies have examined only a limited number of KoV–cell combinations, leaving the host specificity of KoV replication across major vertebrate classes unclear. Here, we systematically compared the replication of mammalian, avian, and reptilian KoVs in cell lines derived from the same three vertebrate classes to characterize the host specificity of KoV replication and examined the relationship between replication competence and KoV phylogeny.

For this comparison, we examined eight KoVs associated with mammalian, avian, or reptilian hosts (Table S1): six mammalian KoVs, HDV, Marmota monax deltavirus (mmDeV), Odocoileus virginianus deltavirus (ovDeV), Desmodus rotundus deltavirus-A (DrDV-A), Desmodus rotundus deltavirus-B (DrDV-B), and Tome’s spiny-rat virus 1 (TSRV-1); one avian KoV, Taeniopygia guttata deltavirus (tgDeV); and one reptilian KoV, Swiss snake colony virus 1 (SwSCV-1). We transfected plasmids encoding 1.2× genome-length copies of each KoV into four mammalian cell lines (Huh-7, U-2 OS, Vero, and Efk3B), three avian cell lines (QT6, LMH, and DF-1), and one reptilian cell line (AOD) (Table S2). We examined the cells at 6, 12, and 18 days post-transfection (dpt) by indirect immunofluorescence assay using antibodies against the delta antigen (DAg) of each KoV and quantified DAg-positive cells as a percentage of DAPI-positive nuclei. We defined KoV replication as persistent when DAg-positive cells remained detectable through 18 dpt (see below discussion and Supplementary Methods for details).

The eight KoVs showed three distinct patterns of host specificity (Figs. 1, 2 and S1). The mammalian KoVs HDV, mmDeV, ovDeV, and DrDV-A replicated persistently in all mammalian cell lines but not in any of the avian or reptilian cells. The avian KoV tgDeV and the reptilian KoV SwSCV-1 showed persistent replication in all avian and reptilian cell lines but not in any of the mammalian cells. In contrast, the mammalian KoVs DrDV-B and TSRV-1 replicated persistently in cell lines from all three vertebrate classes.

**Fig. 1.**
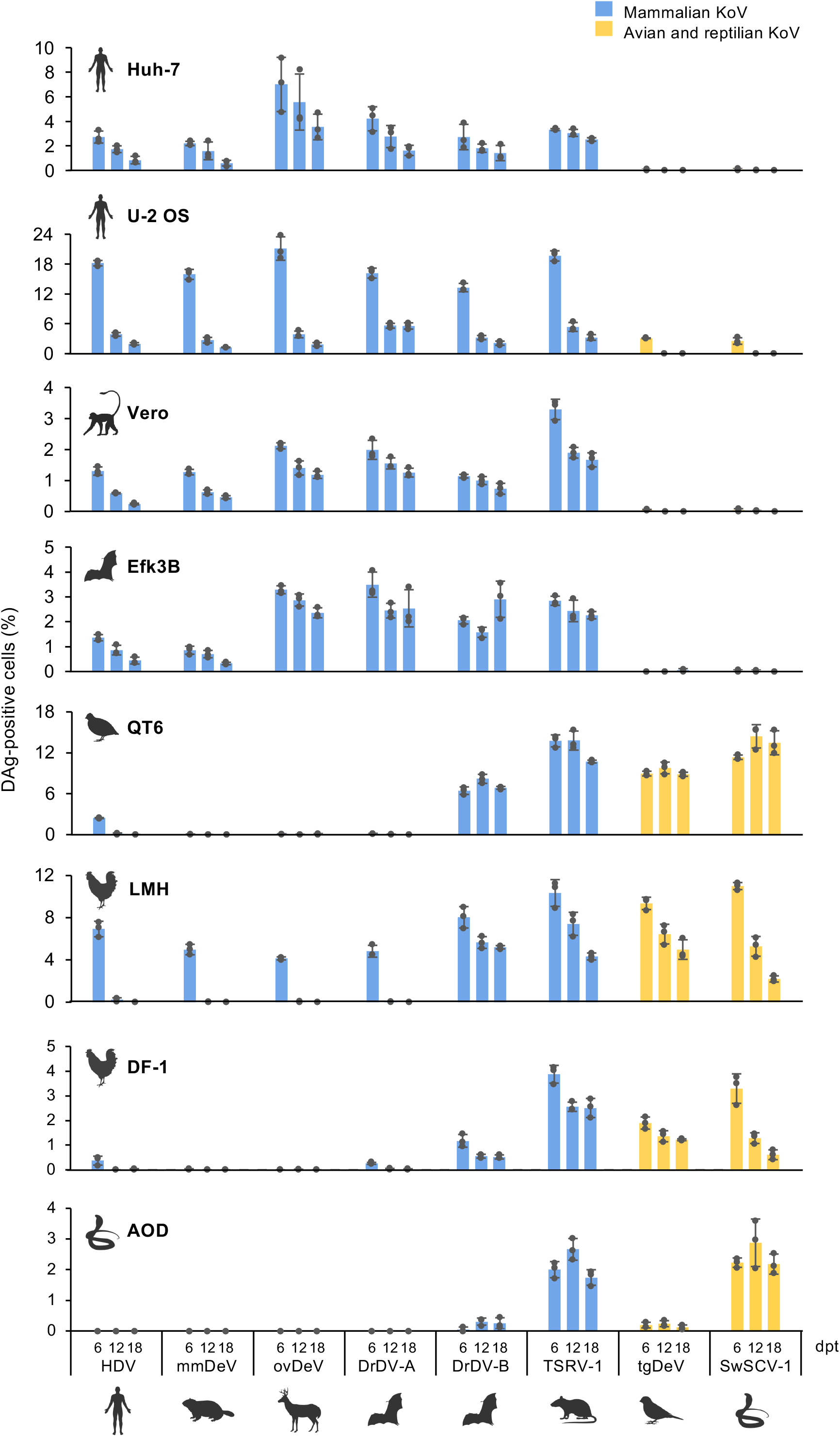
Temporal changes in the proportion of DAg-positive cells in mammalian, avian, and reptilian cell lines. Cells were transfected with plasmids encoding 1.2× genome-length KoV sequences and analyzed at 6, 12, and 18 days post-transfection by indirect immunofluorescence assay using antibodies against the DAg of the corresponding KoV. The percentage of DAg-positive cells was calculated relative to the total number of DAPI-positive nuclei. Data represent the mean ± SD of three independent experiments (n = 3). Representative immunofluorescence images are shown in Fig. S1. Animal silhouettes were obtained from BioRender.com.

**Fig. 2.**
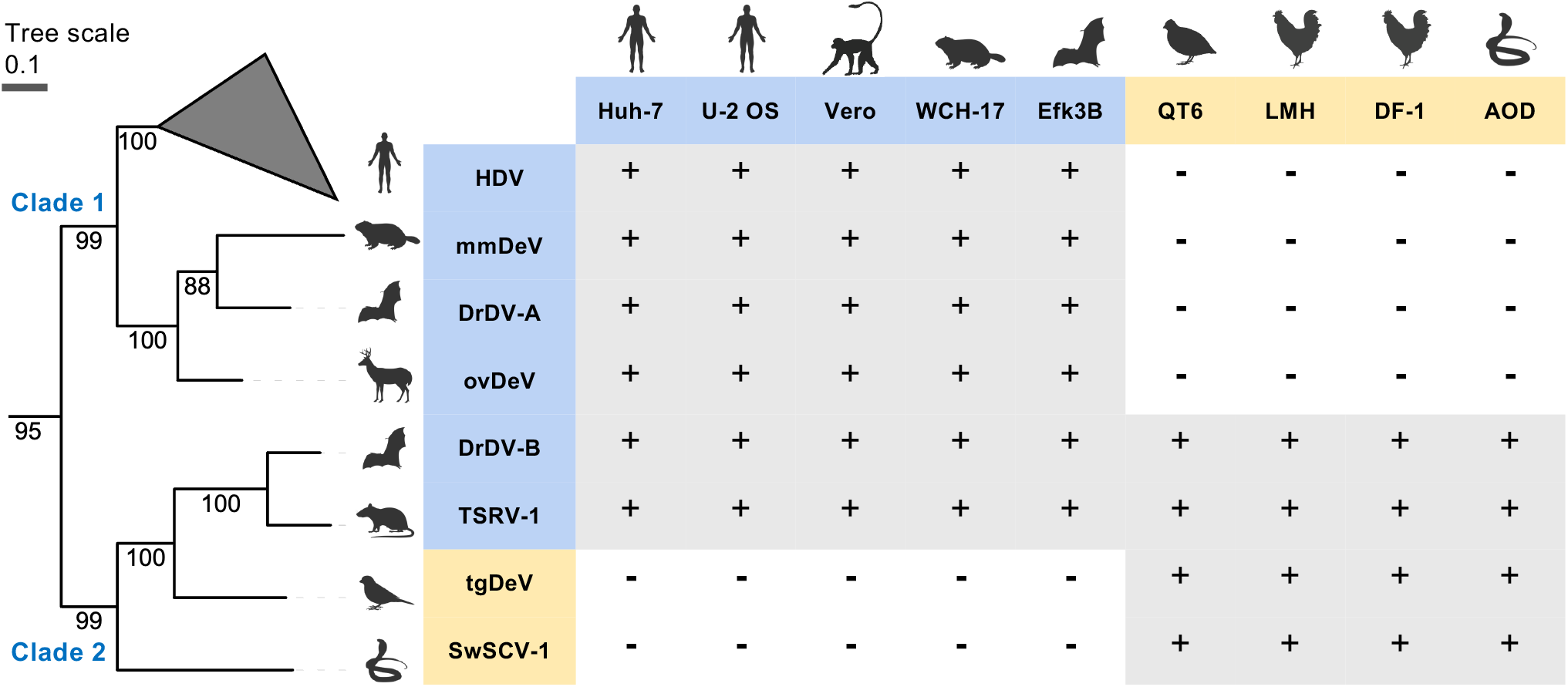
Association between KoV phylogeny and host specificity of persistent KoV replication. A phylogenetic tree based on the DAg amino acid sequences of the KoVs examined in this study is shown together with a table summarizing the persistence of KoV replication in mammalian, avian, and reptilian cell lines. The tree was inferred by the Bayesian Markov chain Monte Carlo method based on an amino acid sequence alignment of DAg. Posterior probabilities are shown for all nodes. The scale bar indicates the number of amino acid substitutions per site. Persistent replication is defined as the presence of DAg-positive cells at 18 days post-transfection (dpt). The presence and absence of DAg-positive cells at 18 dpt are denoted by a plus sign (+) in a gray-shaded cell and a minus sign (−) in an unshaded cell, respectively. Mammalian KoVs and cell lines are highlighted in blue, whereas avian and reptilian KoVs and cell lines are highlighted in yellow. Animal silhouettes were obtained from BioRender.com.

Phylogenetic analysis of the DAg amino acid sequences placed the eight KoVs into two major clades, designated Clade 1 and Clade 2 (Fig. 2). Clade 1 comprises four mammalian KoVs (HDV, mmDeV, ovDeV, and DrDV-A), whose persistent replication was restricted to mammalian cell lines. Clade 2 comprises four KoVs from three different vertebrate classes: the mammalian KoVs DrDV-B and TSRV-1, the avian KoV tgDeV, and the reptilian KoV SwSCV-1. DrDV-B and TSRV-1 are nested within the avian and reptilian KoVs, with SwSCV-1 as the deepest-branching lineage, followed by tgDeV. Notably, the mammalian KoVs within Clade 2 replicated in cell lines from all three vertebrate classes, whereas tgDeV and SwSCV-1 did not replicate in mammalian cell lines. Based on these findings, we propose a scenario for the evolution of the KoVs in Clade 2. In this scenario, the common ancestor of Clade 2 was associated with avian and reptilian hosts. Subsequently, the common ancestor of DrDV-B and TSRV-1 acquired compatibility with mammalian cells while retaining the capacity to replicate in avian and reptilian cells, contributing to the transition of this lineage to mammalian hosts.

Our findings are consistent with previous reports showing that KoVs vary in the range of cell lines in which they can replicate and that some can replicate in cells derived from species other than their known hosts (7, 12, 13). In a comparative study of HDV, TSRV-1, and SwSCV-1 using mammalian and reptilian cell lines, TSRV-1 replicated in both mammalian and reptilian cell lines, SwSCV-1 replicated only in snake cells, and HDV replicated preferentially in mammalian cells (12). The present study extends these observations by comparing a phylogenetically more diverse set of mammalian, avian, and reptilian KoVs in cell lines derived from the same three vertebrate classes, including avian cell lines. Nevertheless, both the phylogenetic diversity of the KoVs and the cells examined remains limited, and extending this approach to additional KoVs and cell lines would provide deeper insights into KoV evolution.

The molecular determinants of the host range of KoV replication remain unknown. KoVs depend on multiple host factors for replication, including host RNA polymerases (14, 15). Previous studies showed that HDV replication was restricted in avian LMH cells but was rescued in avian nuclei after fusion with mammalian cells (16), suggesting that avian cells lack one or more permissive factors required for HDV replication rather than harboring dominant inhibitory factors. The host specificity observed in the present study may likewise reflect differences in the availability or compatibility of host factors required for KoV replication, or alternatively the presence of inhibitory factors in non-permissive cells. Cell fusion experiments such as those used for HDV (16) could distinguish these possibilities. Moreover, the consistent persistence patterns among cell lines derived from the same vertebrate class suggest that at least some of these host requirements are broadly conserved across species and cell types within that class, although only one reptilian cell line was examined.

Our findings also highlight the importance of prolonged observation when assessing persistent KoV replication in cultured cells. For example, in LMH cells transfected with mammalian KoV constructs and in U-2 OS cells transfected with avian or reptilian KoV constructs, DAg-positive cells were detectable at 6 dpt but became undetectable by 12 dpt (Fig. 1). Similar transient DAg expression was previously reported and attributed to transcription driven by a cryptic promoter in the plasmid DNA (12). Alternatively, transient DAg expression could also arise from direct transcription from plasmid-derived KoV RNA, whose highly base-paired structure may permit recognition by host RNA polymerases as a transcription template. Regardless of the underlying mechanism, transient DAg detection alone cannot distinguish such direct expression from transient viral replication. Therefore, assessment beyond one week after transfection is necessary to confirm persistent replication and avoid overestimating the replication host range of KoVs in cultured cells.

This study assessed only intracellular replication following plasmid transfection and did not address helper virus-mediated cell entry, the other major barrier to KoV host specificity. Except for the reptilian KoV SwSCV-1 (11), experimentally validated helper viruses remain unknown for most animal-associated KoVs (17). Future studies identifying helper viruses and incorporating them into experimental systems will be needed to determine their contribution to KoV host specificity and host switching.

Taken together, our findings suggest that changes in intracellular host compatibility contributed to host switching during KoV evolution.

## Acknowledgements

This study was supported by KAKENHI grant numbers 21H01199 (MH), 22K19234 (MH), 23K20902 (MH), 24K21922 (MH), 24K02290 (KW), and 24K18455 (MK), Japan Agency for

Medical Research and Development (AMED) grant number JP25fk0310525 (KW), and the Osaka Metropolitan University (OMU) Strategic Research Promotion Project (Young Researcher) (MK).

