## Supplementary material for "Systematic analysis of intracellular replication competence of mammalian, avian, and reptilian kolmioviruses": Supplmentary Figures

### Fig. S1A

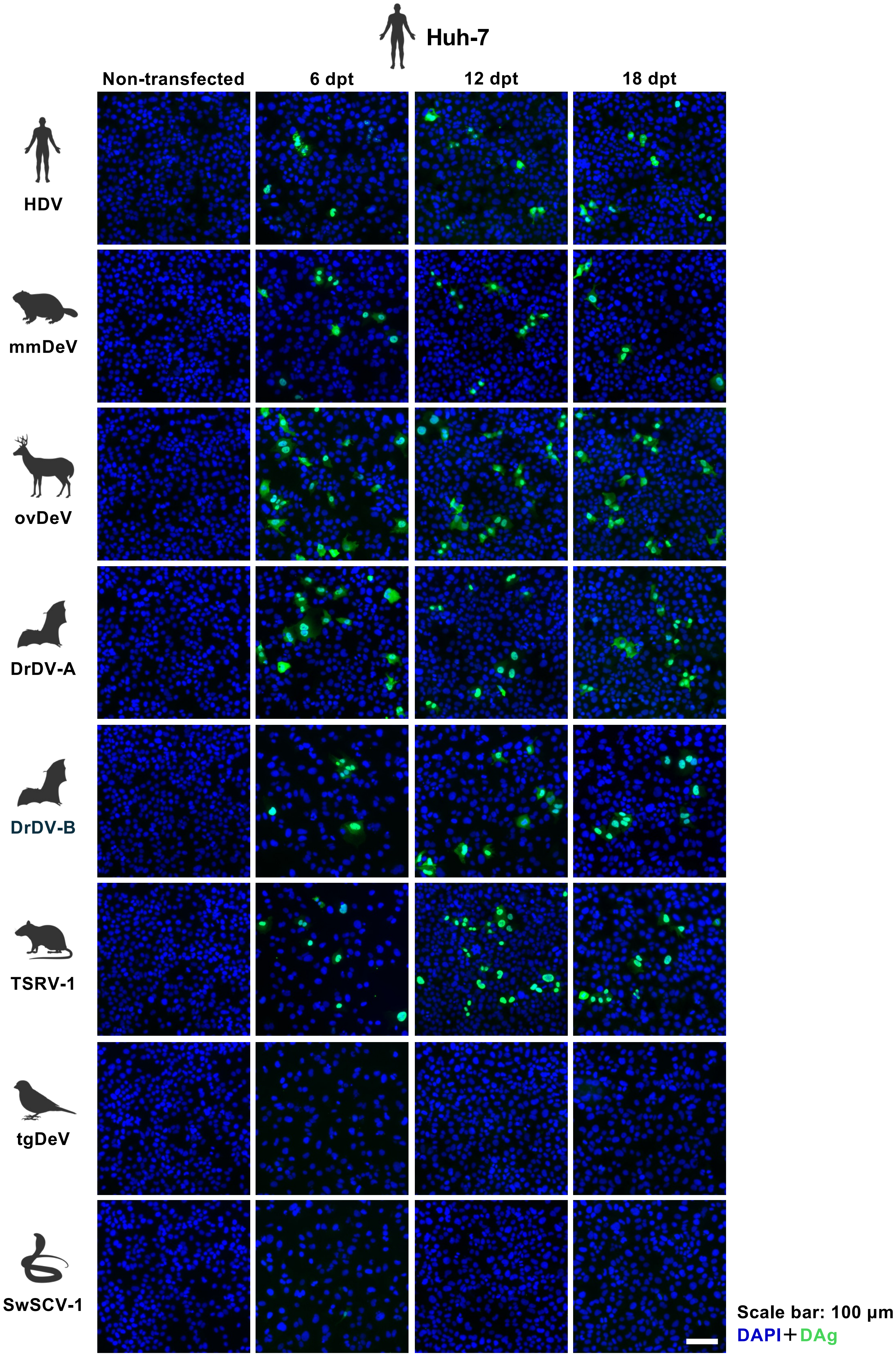

Huh-7
Non-transfected
6 dpt
12 dpt
18 dpt
HDV
mmDeV
ovDeV
DrDV-A
DrDV-B
TSRV-1
tgDeV
SwSCV-1
Scale bar: 100 µm
DAPI+DAg

#### Fig. S1B

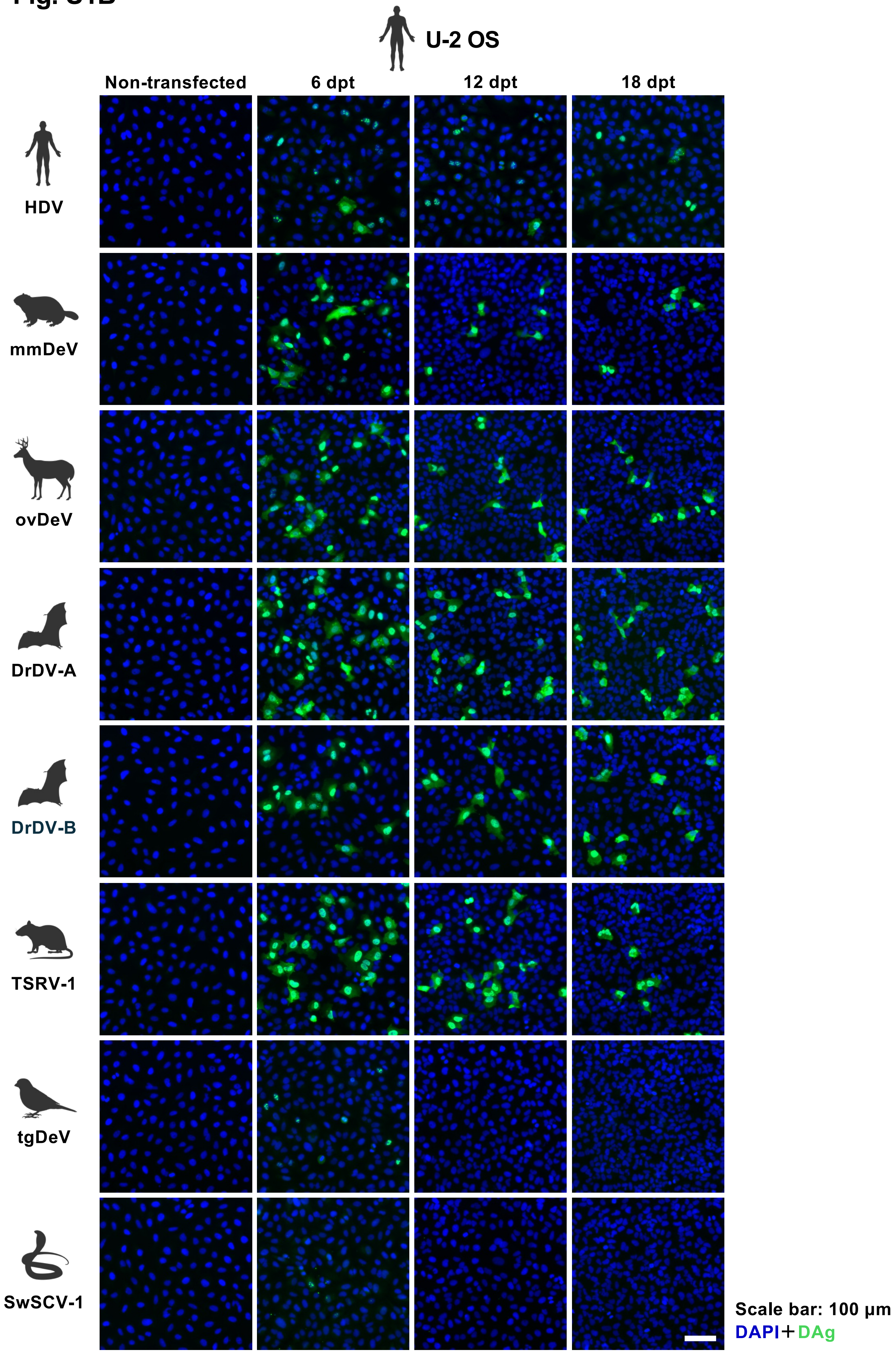
U-2 OS
Non-transfected
6 dpt
12 dpt
18 dpt
HDV
mmDeV
ovDeV
DrDV-A
DrDV-B
TSRV-1
tgDeV
SwSCV-1
Scale bar: 100 µm
DAPI+DAg

Fig. S1C

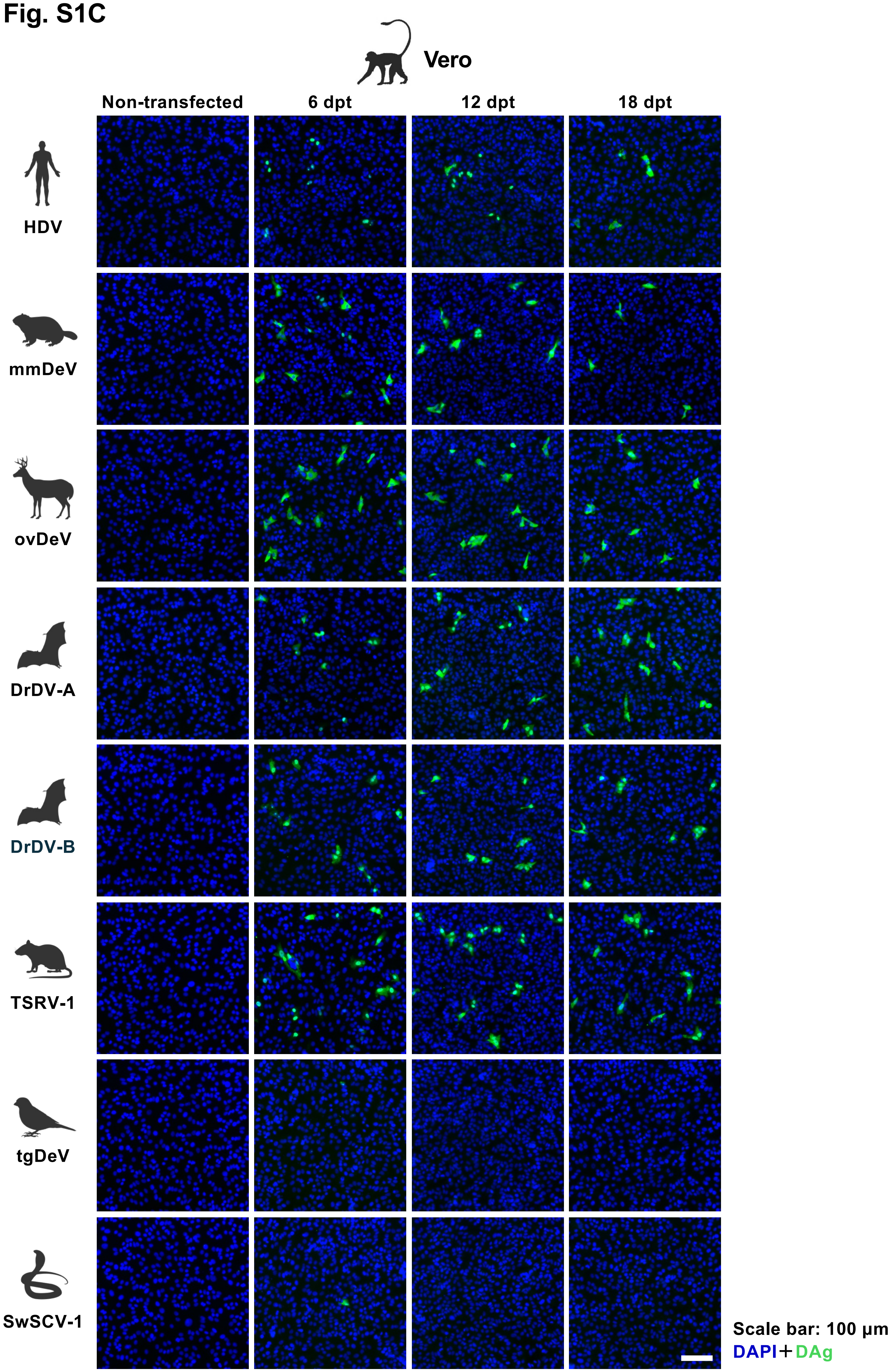
Vero
Non-transfected
6 dpt
12 dpt
18 dpt
HDV
mmDeV
ovDeV
DrDV-A
DrDV-B
TSRV-1
tgDeV
SwSCV-1
Scale bar: 100 μm
DAPI+DAg

Fig. S1D

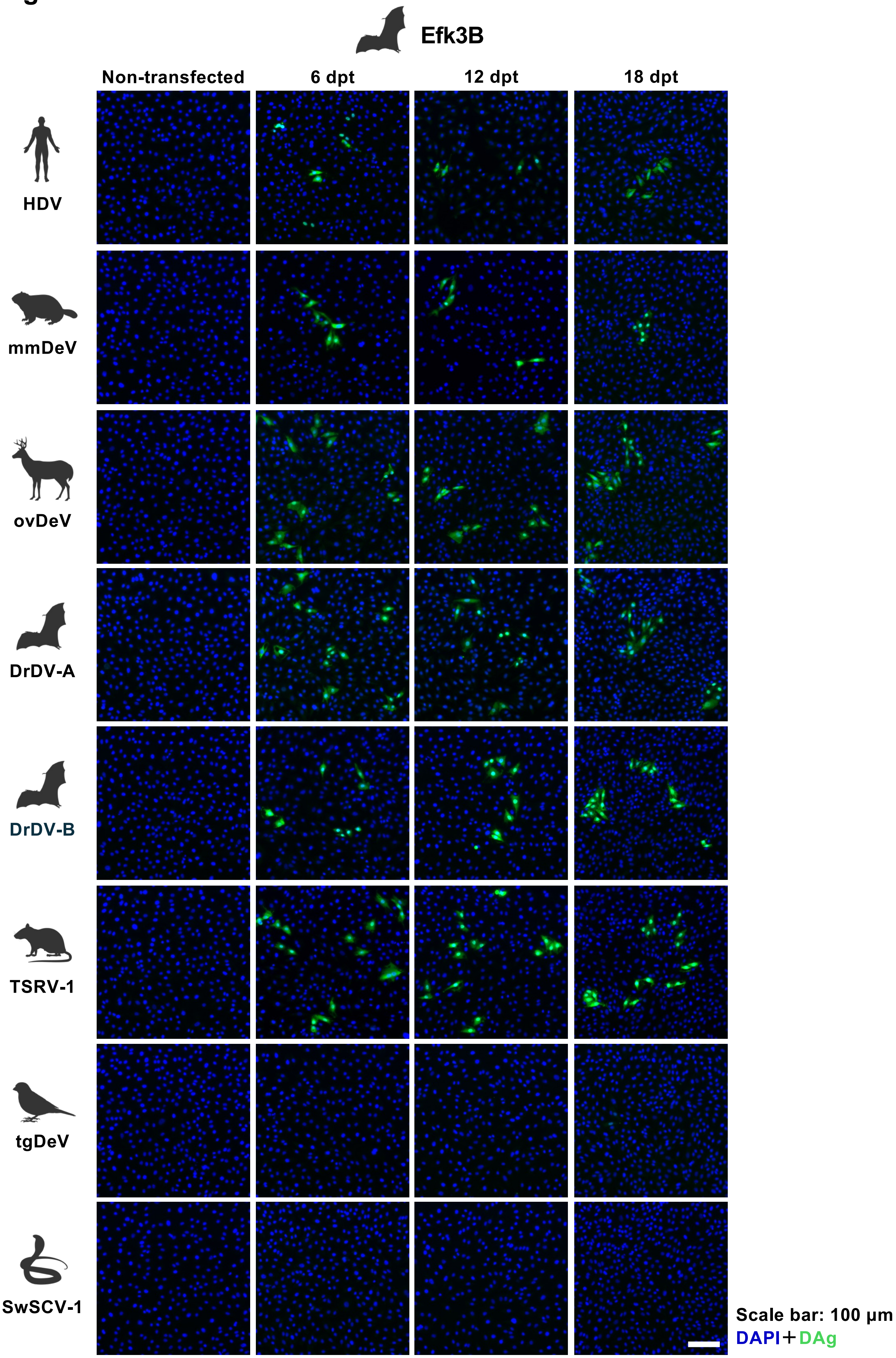

Efk3B
Non-transfected
6 dpt
12 dpt
18 dpt
HDV
mmDeV
ovDeV
DrDV-A
DrDV-B
TSRV-1
tgDeV
SwSCV-1
Scale bar: 100 µm
DAPI+DAg

#### Fig. S1E

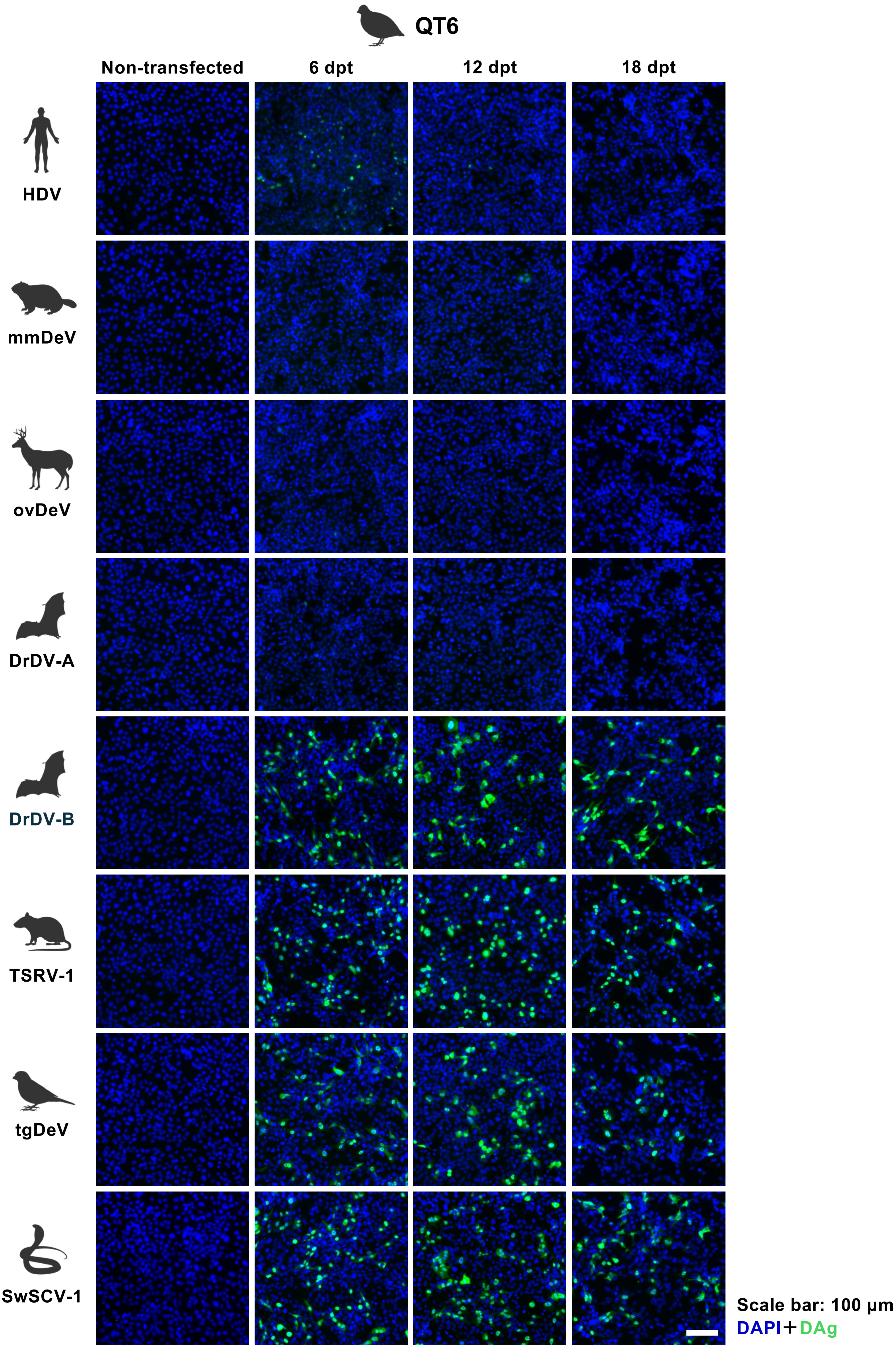

QT6
Non-transfected
6 dpt
12 dpt
18 dpt
HDV
mmDeV
ovDeV
DrDV-A
DrDV-B
TSRV-1
tgDeV
SwSCV-1
Scale bar: 100 μm
DAPI+DAg

#### Fig. S1F

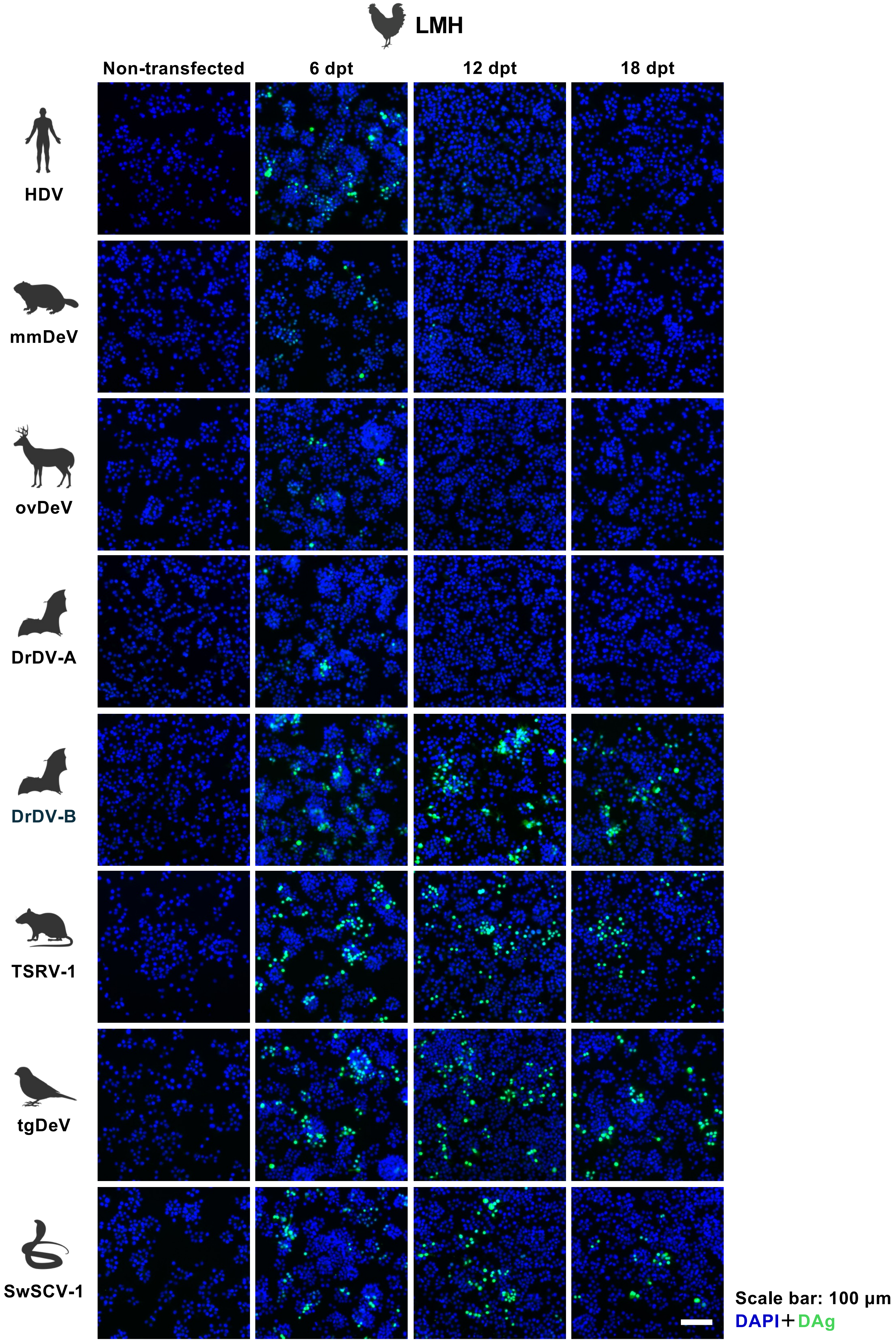
LMH
Non-transfected
6 dpt
12 dpt
18 dpt
HDV
mmDeV
ovDeV
DrDV-A
DrDV-B
TSRV-1
tgDeV
SwSCV-1
Scale bar: 100 μm
DAPI+DAg

#### Fig. S1G

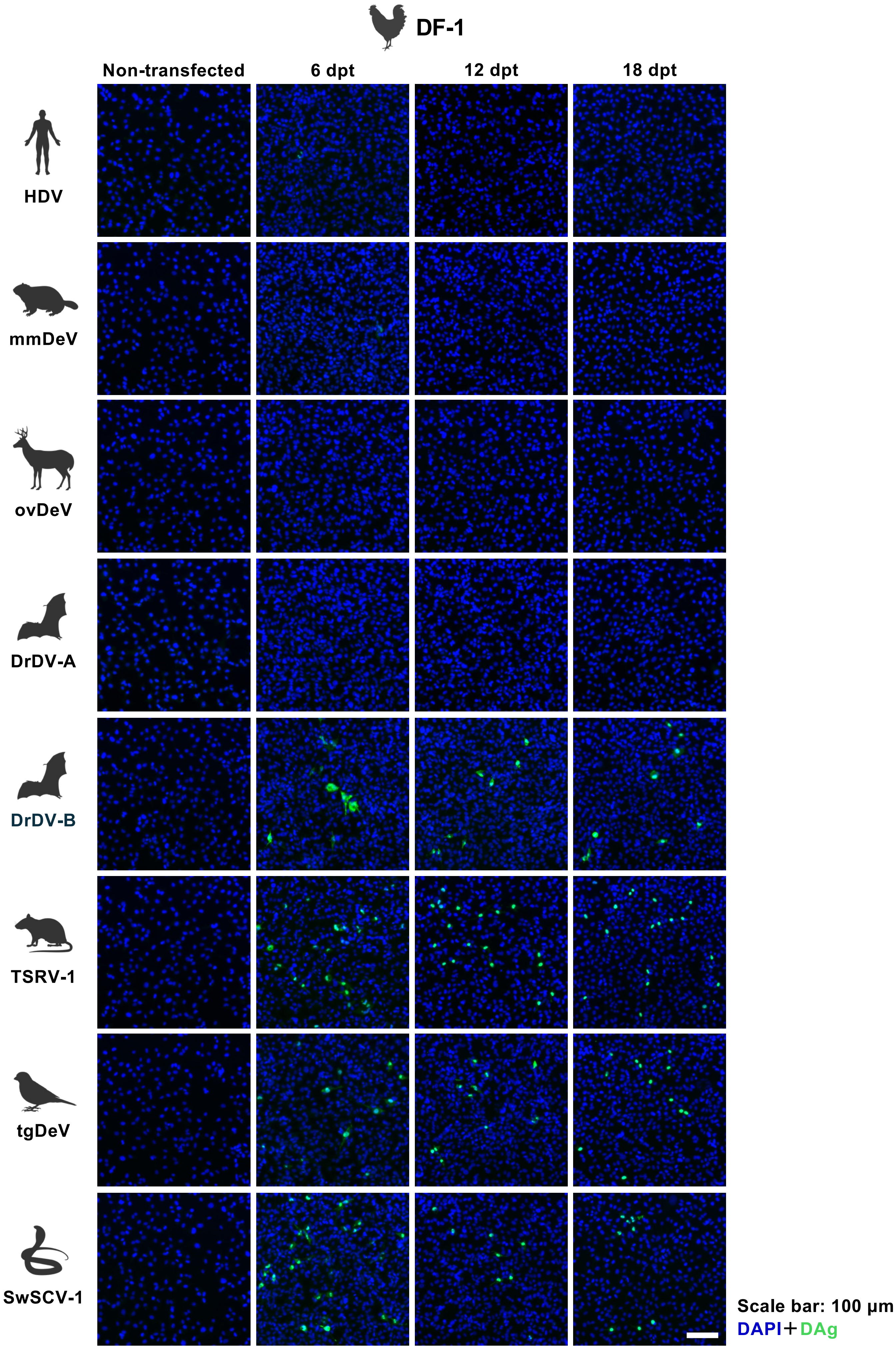
DF-1
Non-transfected
6 dpt
12 dpt
18 dpt
HDV
mmDeV
ovDeV
DrDV-A
DrDV-B
TSRV-1
tgDeV
SwSCV-1
Scale bar: 100 μm
DAPI+DAg

Fig. S1H

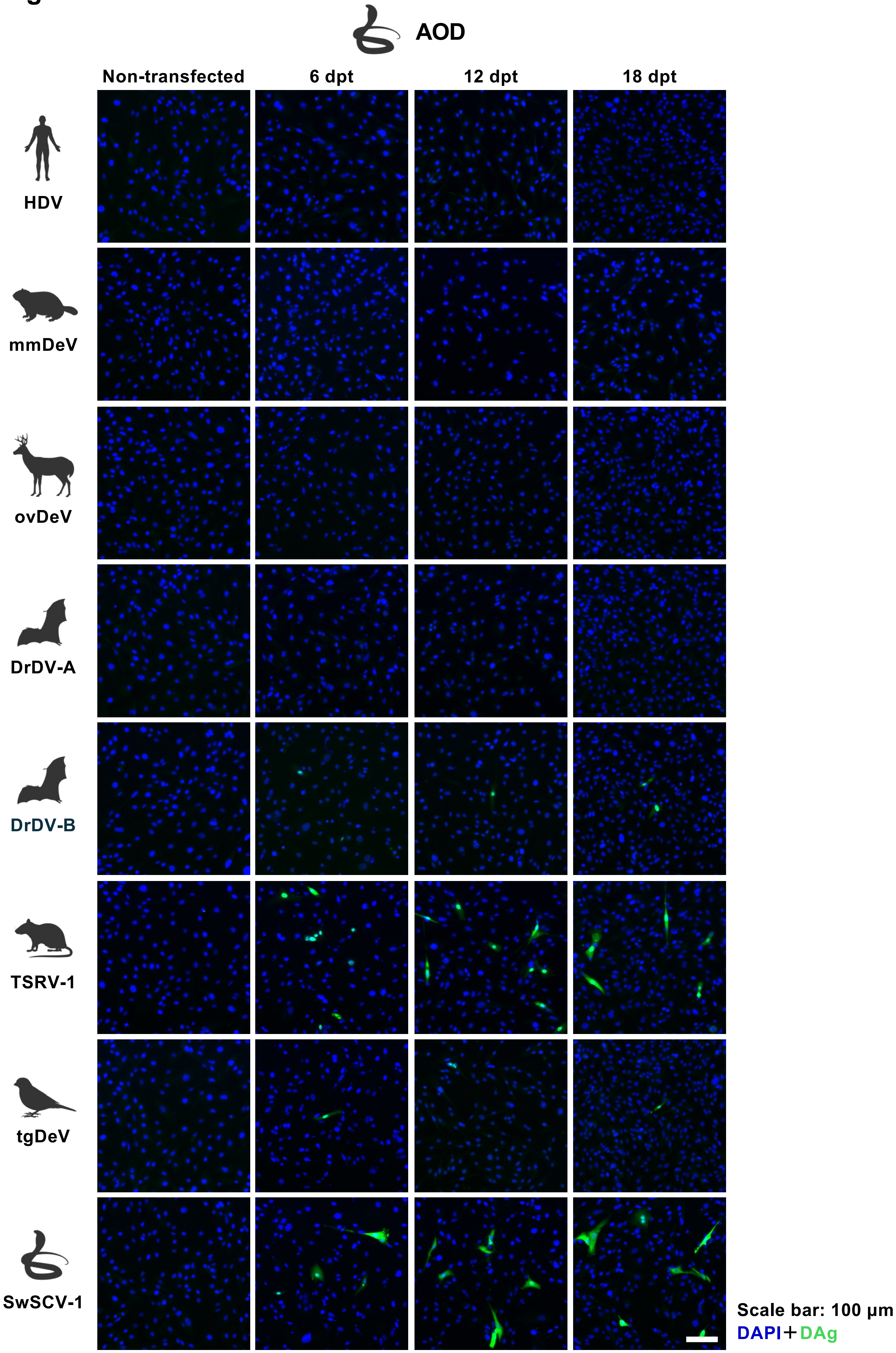
AOD
Non-transfected
6 dpt
12 dpt
18 dpt
HDV
mmDeV
ovDeV
DrDV-A
DrDV-B
TSRV-1
tgDeV
SwSCV-1
Scale bar: 100 μm
DAPI+DAg

**Fig. S1. Representative immunofluorescence images of cells transfected with KoV plasmids over time.**

Representative images of cells ransfected with plasmids encoding 1.2× genome-length KoV sequences at 6, 12, and 18 days post-transfection (dpt) are shown. Non-transfected cells were included as controls. Cells were stained with antibodies against the DAg of the corresponding KoV. (A) Huh-7, (B) U-2 OS, (C) Vero, (D) Efk3B, (E) QT6, (F) LMH, (G) DF-1, and (H) AOD cells. DAg, green; DAPI, blue. Scale bars, 100 μm. Animal silhouettes were obtained from BioRender.com.
