## Supplementary Methods for "Systematic analysis of intracellular replication competence of mammalian, avian, and reptilian kolmioviruses"

### *1. Cell culture*

Huh-7, U-2 OS, Vero, and Efk3B cells were maintained in low-glucose Dulbecco's modified Eagle's medium (DMEM; 08456-36, Nacalai Tesque, Kyoto, Japan) supplemented with 10% fetal bovine serum (FBS; Sigma-Aldrich, St. Louis, MO, USA). DF-1 cells were maintained in the same medium at 39°C. QT6 and LMH cells were maintained in DMEM/Ham's F-12 medium (11581-15, Nacalai Tesque) supplemented with 5% and 10% FBS, respectively. LMH cells were cultured on collagen-coated culture vessels. AOD cells were maintained at 28°C in DMEM supplemented with 15% FBS, 2% non-essential amino acid solution (Nacalai Tesque), 1% MEM vitamin solution (Thermo Fisher Scientific, Waltham, MA, USA), and 1% antibiotic-antimycotic solution (Nacalai Tesque). Unless otherwise indicated, cells were incubated at 37°C in a humidified atmosphere containing 5% $CO_2$. Information on the origins of the cell lines is provided in Supplementary Table 2.

### *2. Plasmids*

Plasmids encoding 1.2× genome-length, negative-sense KoV genomes, spanning from the putative genomic ribozyme to the putative antigenomic ribozyme, were constructed in pTwist-CMV vector. Detailed information on the KoV genomes is provided in Supplementary Table 1.

### *3. Antibody production*

The DAg ORF of each KoV was individually cloned into pET-15b expression vector. *Escherichia coli* Rosetta2(DE3)pLysS cells were transformed with the resulting plasmids and cultured at 37°C in LB medium supplemented with 100 µg/mL ampicillin and 34 µg/mL chloramphenicol. When the cultures reached an $OD_{600}$ of 0.4–0.6, protein expression was induced with 0.4 mM isopropyl β-D-1-thiogalactopyranoside, followed by incubation at 37°C for 3 h. After sonication, recombinant DAg proteins in the soluble fractions were purified using TALON metal affinity resin (Takara Bio, Shiga, Japan). Rabbits were immunized with each purified DAg protein, and the antisera were produced by Eurofins Genomics (Tokyo, Japan). The rabbit antisera were subjected to ammonium sulfate precipitation and dialyzed against phosphate-buffered saline (PBS).

***4. Rescue of KoVs***

The plasmids encoding 1.2× genome-length KoV genomes were transfected into all cell lines except U-2 OS cells using Avalanche-Everyday Transfection Reagent (APRO Science, Tokushima, Japan). U-2 OS cells were transfected using GenJet In Vitro DNA Transfection Reagent for U2OS Cells (SignaGen Laboratories, Frederick, MD, USA). The transfected cells were passaged at a 1:4 split ratio every 3 days, except that AOD cells were passaged at a 1:2 split ratio every 6 days.

***5. Indirect immunofluorescence assay***

Cells were fixed with 4% paraformaldehyde phosphate buffer solution (FUJIFILM Wako, Osaka, Japan) and permeabilized with 0.4% Triton X-100 (Nacalai Tesque) in PBS. After blocking with PBS containing 0.1% Tween 20 and 1% bovine serum albumin (FUJIFILM Wako), the cells were incubated overnight at 4°C with the primary antibodies against the DAg of the corresponding KoV. The cells were then incubated for 1 h at room temperature with Alexa Fluor 488-conjugated secondary antibodies (Thermo Fisher Scientific) and DAPI (FUJIFILM Wako). Fluorescence signals were visualized using an EVOS M7000 fluorescence microscope (Thermo Fisher

Scientific). DAg-positive cells were counted manually, whereas DAPI-positive nuclei were counted using the automated counting function of the EVOS M7000.

#### *6. Phylogenetic analysis*

Phylogenetic relationship among KoVs were inferred as follows. The amino acid sequences of KoV DAg were aligned using MAFFT v7.490 with the E-INS-I algorithm (1). The phylogenetic tree was reconstructed using the Bayesian Markov chain Monte Carlo (MCMC) method in MrBayes v3.2.7a (2) with two independent runs, four chains per run, and the JTT model of substitution. The analysis was run for 5 million iterations, with samples taken every 5,000 steps. The average standard deviation of split frequencies was 0.010. The phylogenetic tree was rooted using seven divergent KoV sequences as outgroups: Chusan Island toad virus 1 (MK962760), Chinese fire belly newt virus 1 (MN031239), Serinus canaria-associated deltavirus (BR001664), dabbling duck virus 1 (MH824555), ray-finned fish virus 1 (MN031240), rhinotermitid virus 1 (MK962759), and Cat Tien Odontotermes delta-like virus (ON082765). The tree was visualized in the Interactive Tree Of Life (iTOL) v6.9.1 (3).
